## Supplementary figures and table for "Distinct sources and behavioral correlates of macaque motor cortical low and high beta"

**SUPPLEMENTARY DISPLAY ITEM LEGENDS**

**Figure S1. Additional behavioral results. Related to Figure 1.**

A. Average hand velocity in one example session in monkey T, split for the three color conditions. On the left, zoomed in to the micro-movements performed during the trial between central touch and GO. To the right with velocity scale adjusted to the final center-out reaching, aligned to movement onset.

B. Hand velocity in a randomly selected subset of correct green trials in the same session as in A. The two solid black vertical lines connected with a horizontal arrow reflect the epoch used to estimate X and Y offset (drift) of micromovements in the post-cue epoch (in D). The velocity scale is indicated inside the plot.

C. Average hand velocity in the 1s delay after each SC, split for color condition, averaged across all trials for all behavioral sessions for the two monkeys combined. Horizontal black lines on top of the bar plots denote significant differences in single-trial hand velocity.

D. Hand cursor displacement (drift) caused by micro-movements across all green trials in each monkey, split according to the target direction. Each dot reflects one trial, and the position reflects the relative X and Y offset 1s after the onset of SC2 (2nd vertical solid black line in B), compared to the position at SC2 onset (1st vertical solid black line in B). UR-upper right; LR-lower right; LL-lower left; UL-upper left. The total number of trials is indicated (n).

E. Average deltoid EMG amplitude, recorded in the same behavioral session as shown in A-B, on the left for the period between touch and GO (split for the three color conditions) and on the right aligned to movement onset (averaged for the three color conditions). Towards the body (LL) in solid lines and away from the body (UL) in dotted lines. The raw EMG signal (30kHz) was first rectified, then low-pass filtered at 250Hz and downsampled to 1kHz. A Gaussian filter (length 150ms, width 100ms) was used to smooth single trials before plotting the trial-averaged EMG.

**Table S1. Behavioral task performance. Related to Figure 1.**

Summary of all errors, number of correct trials included for behavioral analyses, percent of distractor errors and RTs for each color condition and movement direction for each animal. UR-upper right; LR-lower right; LL-lower left; UL-upper left.

**Figure S2. Average normalized power and peak frequency distribution of the full signal. Related to Figure 3.**

A. Average normalized power in the pre-SC1 period across all trials for all sites in each monkey, for the full LFP signal, including aperiodic and periodic components. The curves reflect the mean power ±SEM across LFP sites. Overlain are distributions of single-trial peak frequency (frequency with maximal power) between 10-40Hz in the same task period for the full LFP signal.

**Figure S3. Average low and high beta amplitude, and main regressors explaining amplitude variance. Related to Figure 4.**

Same as Figure 4, separated for Monkey T (A,B) and Monkey M (C,D).

A,C. Representation of the trial-averaged temporal profile of normalized high (left ; 21-29Hz) and low (right ; 12-20Hz) beta amplitude (+/-SEM), separated by the 3 color conditions. The horizontal gray line above each plot graph represents the time-resolved modulation in beta amplitude by the color condition along the task. The significativity is represented as in Figure 4A.

B,D. Time-resolved representation of the presence of each regressor in the winning model after the application of a Bayesian Index Criterion (BIC) for the comparison of all possible models and their 2-by-2 interactions, for the high beta (left) and the low beta (right). Each row represents a regressor, the last row represents all possible interactions. Regressors selected for ulterior analysis in orange and discarded regressors in gray.

**Figure S4. Average MUA in high and low beta dominant sites. Related to Figure 4.**

Average MUA amplitude including all trials of all recording sites, for high beta band (left) and low band (right) dominant sites, separately for the three color conditions. The MUA was generated following the method of Stark and Abeles[1]. The raw signal was first bandpass filtered (300-6000 Hz) and clipped beyond +/-2 standard deviations. Then, the signal was squared, smoothed with a low-pass filter (250Hz) and downsampled from 30 to 1 kHz, before the square-root was taken to arrive at the final MUA signal. The MUA from each site was cut in trials, and normalized by dividing by the mean amplitude across all trials and trial-times. Finally the single-trial MUA from all sites with the same LFP beta band dominance were combined. The plots show trial-averaged MUA after first smoothing individual trials with a Gaussian filter (length 30ms, width 15ms).

**Figure S5. Variance explained by full model with each regressor scrambled across trials. Related to Figure 4.** A. Total percentage of variance explained by the 4 selected regressors, RT, hand Velocity, gaze position and time-on-task, along the task, in the 3 different color conditions for both bands. B-E. Percentage of variance explained by the full model minus the full model in which the values of one regressor were scrambled across trials. For each bin the values were scrambled 100 times and the average of the 100 scrambles was subtracted to values obtained with the full model. The dots above each graph represents an equivalent p-value of 0.01. For a temporal bin, if the value of variance explained obtained with the full model was superior to all the 100 values obtained with the regressor of interest scrambled, the effect of the regressor in that bin was considered significant and marked with a dot. From B to E, the values were scrambled respectively for the RT, hand Velocity, gaze position and time-on-task.

**Figure S6. Influence of RT on low beta depending on the presence of other regressors. Related to Figure 5.**

Comparison of the significativity of RT as the unique regressor of a linear model vs. paired with each of the other regressors, separated by color conditions. In each plot, the p-value are displayed on a logarithmic scale, in color when RT was the unique regressor considered and in black when paired with a second regressor. The horizontal dotted lines on the top of each plot represent the significance of each case ($p<0.01$). Red horizontal dotted lines are plotted for p-values equal to 1, 0.05 and 0.01. A. RT paired with hand velocity as a second regressor. B. RT paired with the gaze position as second regressor. C. RT paired with time-on-task as second regressor.

**Figure S7. Correlations between beta amplitude and gaze position. Related to Figure 7.**

A. Representation of the negative, zero and positive lag meaning in the correspondence between LFP (top) and gaze position (bottom). Three trials are represented for beta and gaze position. A negative lag means relating the gaze position with LFP in the past (before), beta was consequently leading gaze. Zero lag means relating LFP and gaze position from the same temporal bin. A positive lag means relating gaze position to the LFP in the future (later), gaze was consequently leading beta. B. Proportion of bins in which beta amplitude (both bands combined) modulated significantly with gaze position, for different temporal lags. Beta was leading gaze for negative values (i.e. gaze at time $t0$ and LFP at time $t0$ - lag, represented in orange). Correlation at zero lag is represented in gray. Gaze was leading beta for positive values (i.e. gaze at time $t0$ and LFP at time $t0$ + lag, represented in red). C. Low beta band split into groups of trials based on the position of the gaze at 200ms after the onset of the valid SC. Brown curves represent the trials in which the monkeys were looking inside the working area (either on the target, or elsewhere). Purple represents the trials in which the monkeys were looking outside the working area. D. High beta amplitude split into groups based on the monkey's position of the gaze at different time lags (From top to bottom. -1000 ms, -240 ms, 0 ms, 240 ms, 1000), all color conditions combined. Red dashed vertical lines represent the values from and up to which significant bins have been counted in B. We excluded the first and last 1000ms of the period spanning from -1200ms to 5600ms from SEL. E. Same representation for the low beta band.

**Figure S8. Spectral parametrization for early and late trials. Related to Figure 8.**

A. Spectral parameterization using the FOOOF method[2] for the pre-SC1 period of blue trials. The analysis was done separately for the first third of trials for each session (early; left) and the last third of trials for each session late; right), for each monkey separately. The black line corresponds to the original data and the red line to the model fit. The algorithm identifies the aperiodic signal (blue dashed line) and the spectral peaks and their peak frequency (green). A frequency range of 5-194Hz was used for fitting the data, using the ‘knee’ mode.

B. Spectrum decomposition in periodic (left) and aperiodic (right) signal components, in early and late blue trials in the sessions, for each monkey. The frequency axis was cut at 45Hz for the periodic signal to focus on the lower frequencies including the beta bands.

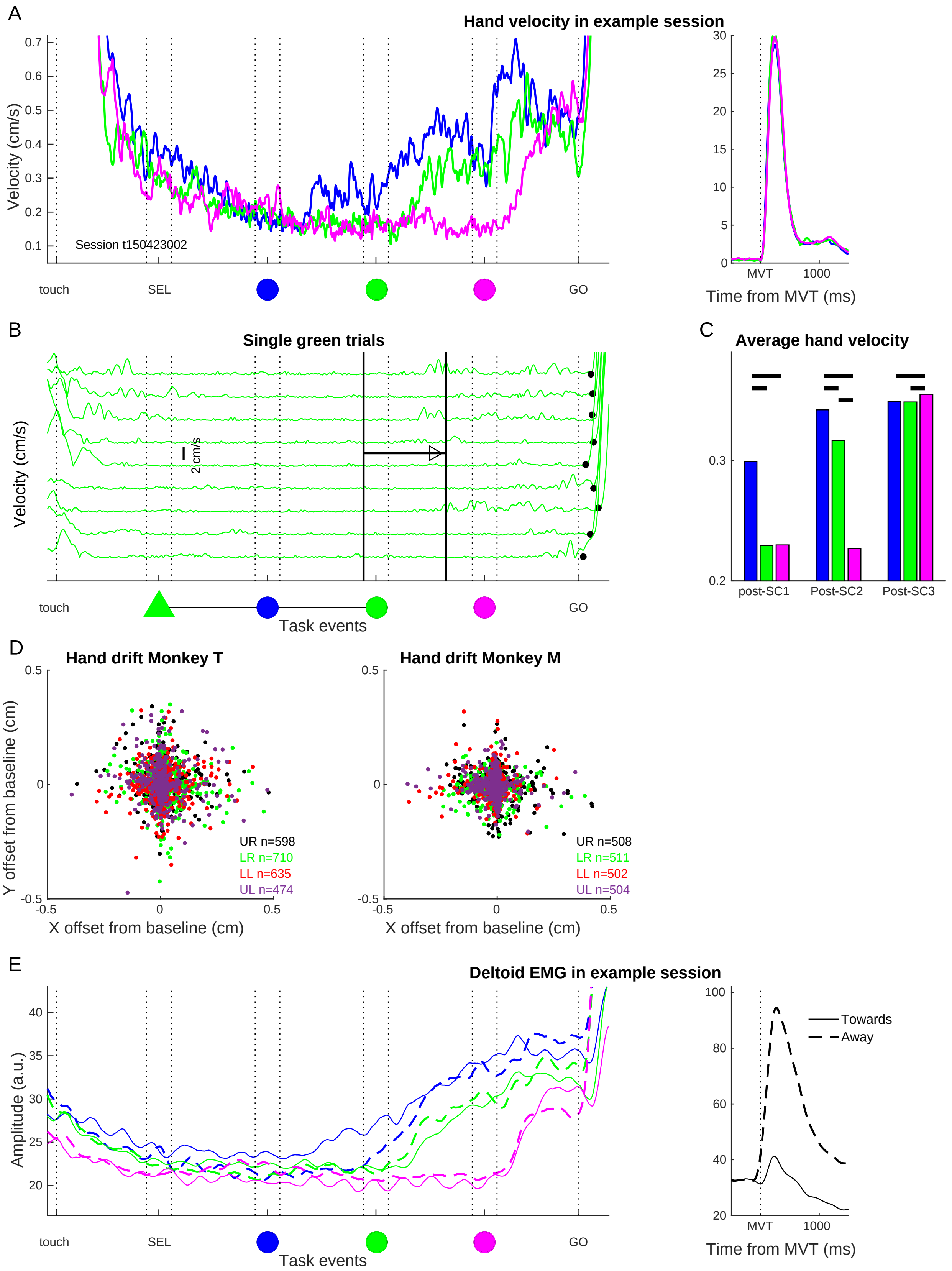

A
Hand velocity in example session
Velocity (cm/s)
Session t150423002
touch
SEL
GO
Time from MVT (ms)
MVT
1000
B
Single green trials
Velocity (cm/s)
2 cm/s
Task events
C
Average hand velocity
post-SC1
Post-SC2
Post-SC3
D
Hand drift Monkey T
Hand drift Monkey M
Y offset from baseline (cm)
X offset from baseline (cm)
UR n=598
LR n=710
LL n=635
UL n=474
UR n=508
LR n=511
LL n=502
UL n=504
E
Deltoid EMG in example session
Amplitude (a.u.)
Towards
Away


Figure S1

**Overview of all movement errors**

| | Abort pre-GO (% of initiated) | Long RT (% of GO trials) | Long MVT (% of GO trials) | Touch uncued (% of GO trials) | Touch distractor (% of GO trials) |
|---|---|---|---|---|---|
| **Monkey T** | 41.9 | 1.3 | 7.4 | 1.7 | 17.4 |
| **Monkey M** | 37.8 | 4.4 | 2.2 | 3.0 | 22.3 |

**Number of correct behavioral trials for each color condition and movement direction**

| | Blue | Green | Pink | UR | LR | LL | UL |
|---|---|---|---|---|---|---|---|
| **Monkey T** | 2261 | 1987 | 1766 | 1556 | 1647 | 1577 | 1234 |
| **Monkey M** | 2109 | 1821 | 1643 | 1396 | 1406 | 1383 | 1388 |
| **Proportion of distractor errors (% of correct + distractor)** | | | | | | | |
| **Monkey T** | 20.7 | 20.4 | 16.7 | 19.4 | 16.8 | 18.6 | 23.7 |
| **Monkey M** | 30.8 | 27.3 | 11.2 | 27.1 | 23.5 | 23.9 | 24.2 |
| **Reaction times in correct trials, from hand trajectories (ms)** | | | | | | | |
| **Monkey T** | 150 +/-40 | 152 +/-39 | 147 +/-50 | 150 +/-50 | 150 +/-38 | 147 +/ 37 | 152 +/-46 |
| **Monkey M** | 173 +/-57 | 169 +/-58 | 153 +/-65 | 170 +/-57 | 150 +/-62 | 163 +/-60 | 179 +/-59 |

**Table S1**. Summary of all errors, number of correct trials included for behavioral analyses, percent of distractor errors and RTs (+/-SD) for each color condition and movement direction for each animal. UR-upper right; LR-lower right; LL-lower left; UL-upper left.

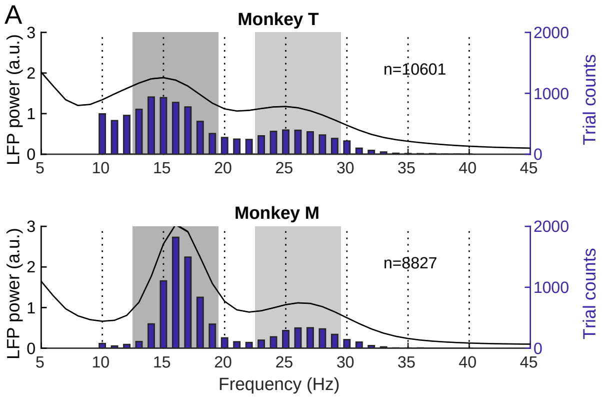
A
Monkey T
LFP power (a.u.)
Trial counts
n=10601
Monkey M
n=8827
Frequency (Hz)

Figure S2

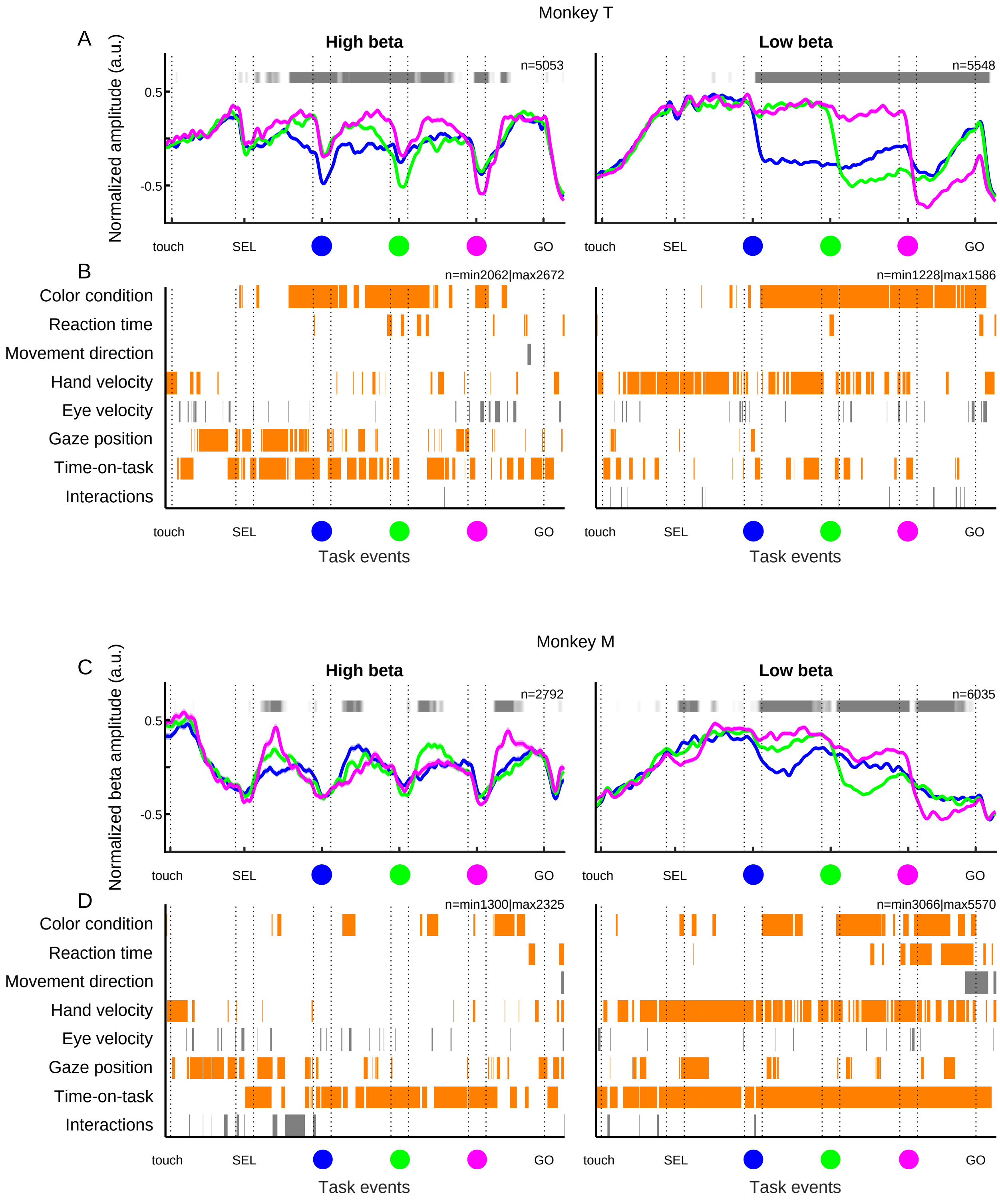
Monkey T
A
High beta
Low beta
n=5053
n=5548
0.5
-0.5
Normalized amplitude (a.u.)
touch
SEL
GO
B
n=min2062|max2672
n=min1228|max1586
Color condition
Reaction time
Movement direction
Hand velocity
Eye velocity
Gaze position
Time-on-task
Interactions
Task events
Monkey M
C
n=2792
n=6035
Normalized beta amplitude (a.u.)
D
n=min1300|max2325
n=min3066|max5570

Figure S3

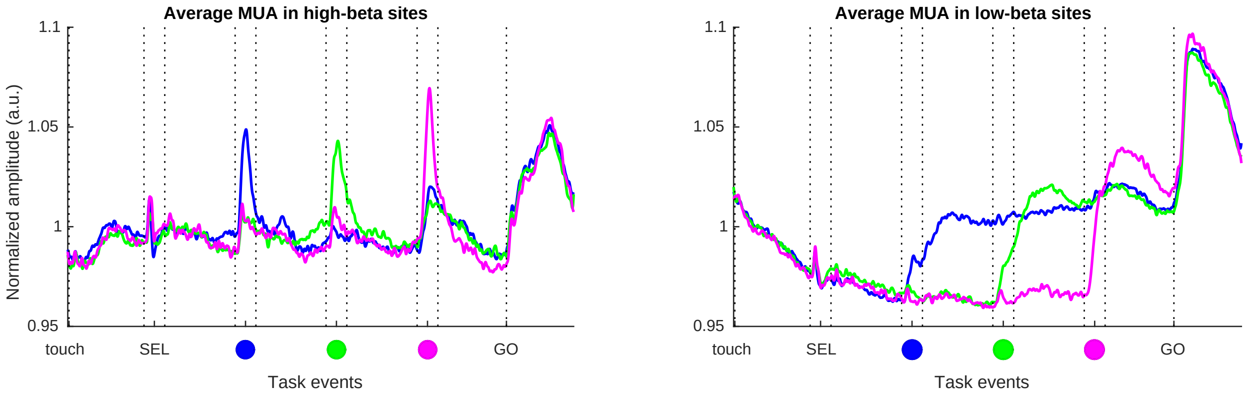
Average MUA in high-beta sites
Average MUA in low-beta sites
Normalized amplitude (a.u.)
1.1
1.05
1
0.95
touch
SEL
GO
Task events

Figure S4

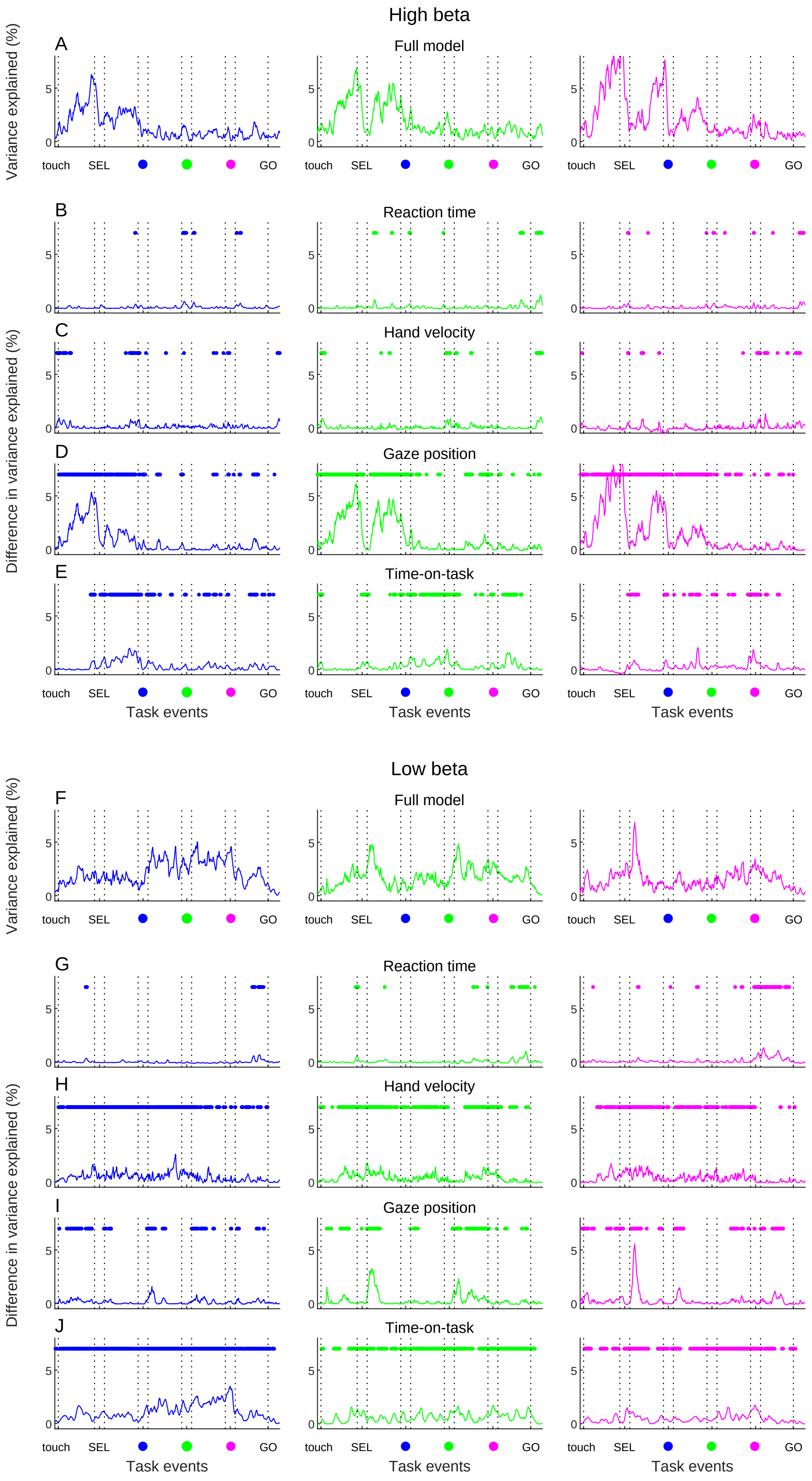
High beta
Full model
Reaction time
Hand velocity
Gaze position
Time-on-task
A
B
C
D
E
Variance explained (%)
Difference in variance explained (%)
touch
SEL
GO
Task events
Low beta
F
G
H
I
J

Figure S5

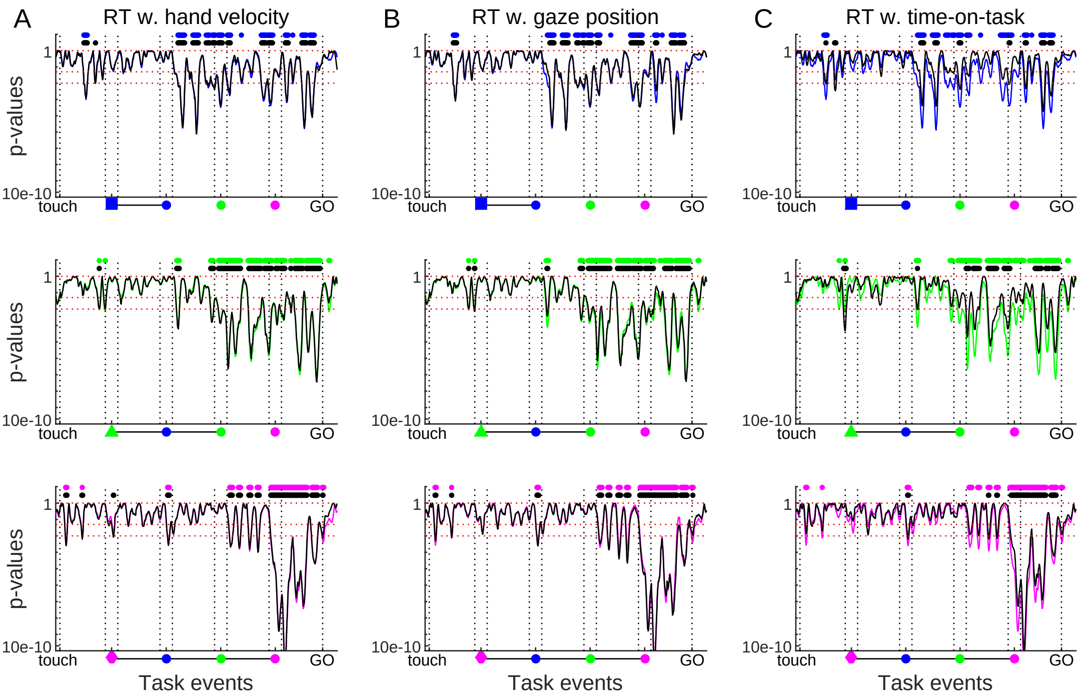

A
RT w. hand velocity
B
RT w. gaze position
C
RT w. time-on-task
p-values
1
10e-10
touch
GO
Task events


Figure S6

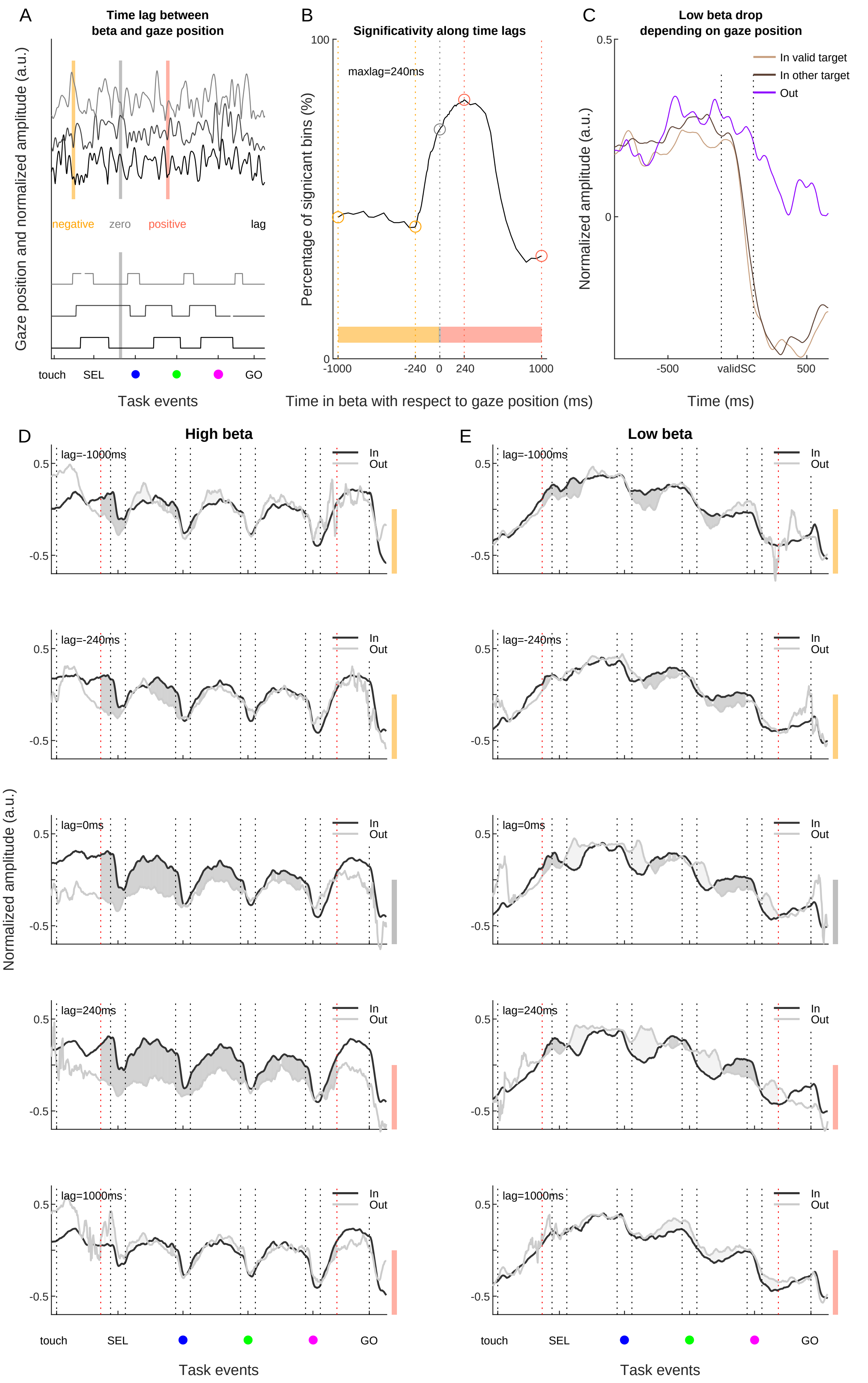
A
Time lag between
beta and gaze position
Gaze position and normalized amplitude (a.u.)
negative
zero
positive
lag
touch
SEL
GO
Task events
B
Significativity along time lags
maxlag=240ms
Percentage of signicant bins (%)
100
0
-1000
-240
0
240
1000
Time in beta with respect to gaze position (ms)
C
Low beta drop
depending on gaze position
In valid target
In other target
Out
Normalized amplitude (a.u.)
0.5
0
-500
validSC
500
Time (ms)
D
High beta
E
Low beta
lag=-1000ms
lag=-240ms
lag=0ms
lag=240ms
lag=1000ms
In
Out
0.5
-0.5
Normalized amplitude (a.u.)
touch
SEL
GO
Task events

Figure S7

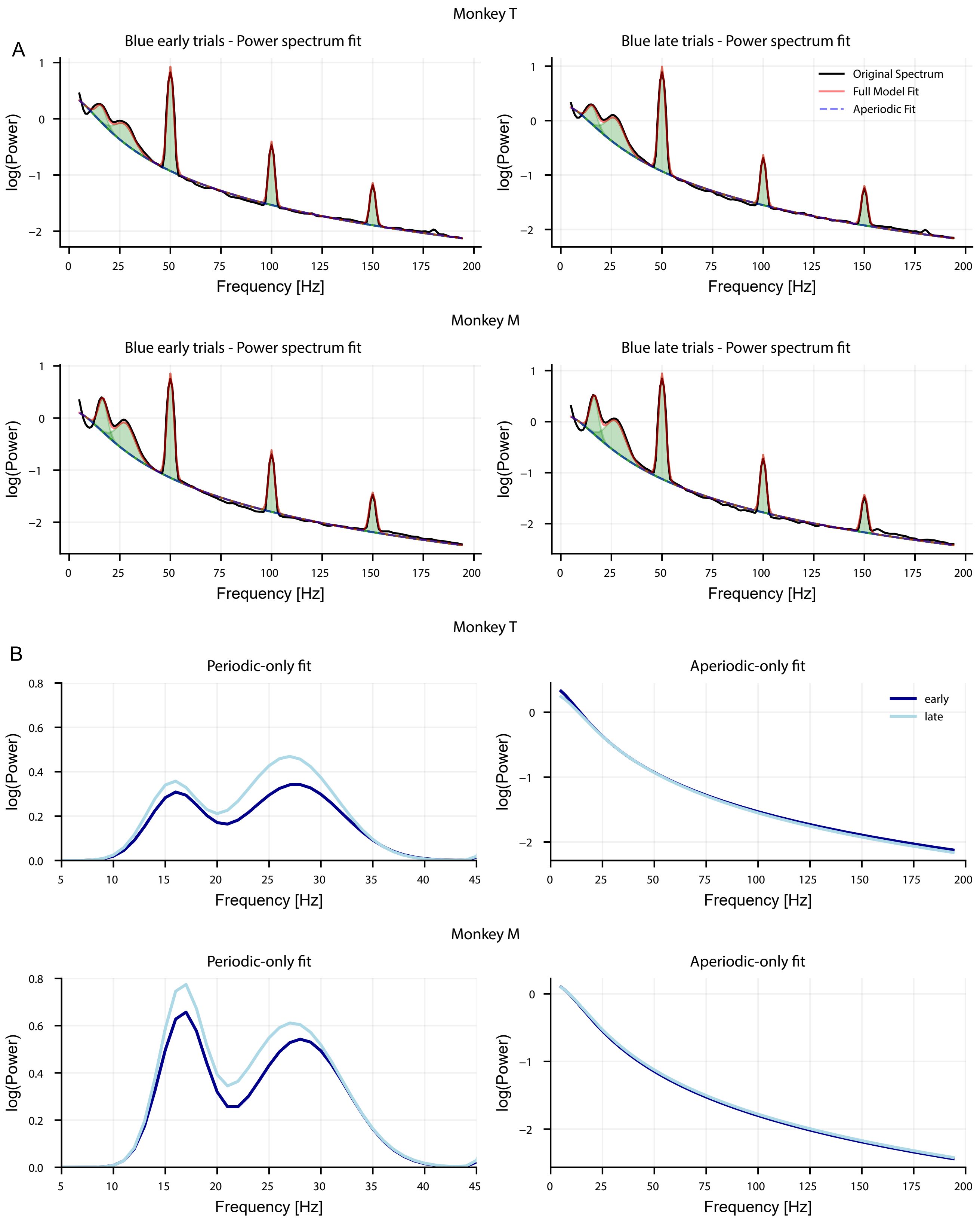
Monkey T
A
Blue early trials - Power spectrum fit
Blue late trials - Power spectrum fit
Original Spectrum
Full Model Fit
Aperiodic Fit
log(Power)
Frequency [Hz]
Monkey M
Blue early trials - Power spectrum fit
Blue late trials - Power spectrum fit
Monkey T
B
Periodic-only fit
Aperiodic-only fit
early
late
Monkey M
Periodic-only fit
Aperiodic-only fit

Figure S8
